# Magnetocontrolled protein membranes for cell cultures co-cultivation

**DOI:** 10.1101/2020.06.05.135897

**Authors:** A. Minin, M. Tiuchai, S. Rodionov, I. Blatov, I. Zubarev

## Abstract

Investigation of cells and tissues in vitro systems is an attempt to simplify the very complex interactions between the various cell types from multicellular organisms. Monolayer cell cultures with single cell type do not allow to show all the possible paracrine interactions between various types of cells. To analyze intercellular interactions, it is possible to use systems that co-cultivate several cell types. This article proposes a new cell co-cultivation system based on levitation in the magnetic field in the culture medium of a magnetic protein membrane with cells. The developed system of co-cultivation of cells can be made in any laboratory of available reagents and have a low manufacturing cost.

## 1 INTRODUCTION

The ability to reproduce in vitro cell-cell interactions and transition from one functional state to another provides researchers with model systems for studying various physiological processes and pathological states. In a single cell type culture, the unique three-dimensional tissue structure is lost, while in a combined culture, the availability of direct cell-cell contacts can influence the functional state of the cells and it is impossible to evaluate only the paracrine effect of the cells on each other. In such cultures, it is not possible for individual cell populations and study them separately. Withal, the separation of cell types in the culture vessel allows evaluating only their paracrine effect. These molecular interactions can determine cell metabolism, which allows estimating the effect of co-cultivation on the growth and physiological state of one or both cell types without labeling of the cell population.

There are several types of co-cultivation systems with various cell lines that are applied to study the paracrine effect between cells and modeling the in vivo processes. The simplest model of cell co-cultivation is to apply a conditioned medium: two cell populations are cultivated separately, then the culture medium from one population is collected and used for another cell population. Withal, short-lived molecules are not stable in a conditioned medium and do not have time to transfer to the wells with a second cell population. In addition, most long-lived molecules will be diluted over time through diffusion. In these cases, both cell populations interact only in unidirectional paracrine communication that leads to the absence of feedback signals existing in natural conditions ^1^. It was proposed to grow cells by mechanically limiting cell growth in the center of the Petri dish using a metal rod followed by seeding the free area with another cell type ^2^.

One of the most significant achievements was the use of permeable inserts with microporous membranes for the compartmentalization of cell cultures, first used by Grobstein in 1953^3^. The authors proposed a variant of indirect cell culture using sterile disposable inserts with different sizes, which are inserted into cultured Petri dishes or wells of a multiwell plate in the culture medium. Such systems for modeling changes in the cellular phenotype are called the “Transwell system” (Corning Costar Corp., Cambridge, MA, USA). They present a semipermeable membrane, allowing the cultivation of two different types of cells, sharing the common culture medium, but without direct cell-cell contact. In the co-cultivation system, the insert contains one cell type, while the bottom of the culture vessel contains another cell population, which allows studying the contribution of two different cell populations to their total biochemical environment. As a result, cell polarity is maintained, which gives co-culture systems an important advantage over the methods of mixed cultures and conditioned medium. The limitation of such systems is the lack of real-time tracking of adhesion/transmigration events and control of the biochemical/biophysical parameters of the local microenvironment.

In such a system, paracrine signaling occurs between cells, without direct intercellular contact. The secretion of proteins, such as growth factors and cytokines, affects cell behavior, proliferation, maturation and differentiation ^4^. These effector molecules, which are produced by a wide range of cells, including immune cells, such as macrophages and lymphocytes, as well as fibroblasts and endothelial cells (ECs), are critical for activating the immune system through various receptors ^5^.

Several experiments have shown that indirect co-cultivation systems are effective for producing a sufficient amount of cytokines. Inserts were used in several papers in the inflammation research field. The effects of secreted cytokines in various immune cells and processes of inflammation in the central nervous system (CNS) were assessed ^6,7,8,9,10,11^. In the co-culture systems for CNS research stem cells are used to study neurotrophism and neuroprotection ^12,13^. These systems have also been developed to study the anti-inflammatory potential of molecules, which depends on their ability to reduce or inhibit the secretion of pro-inflammatory factors ^14,15^. Additionally, membrane inserts can be used during cancer studies ^16,17^ to understand the mechanisms underlying angiogenesis ^18^, inflammation ^19,20^ in oncogenesis and process of cell differentiation ^21,22^. Also, Transwell system inserts are used in such diverse areas as nephrology ^23^, endothelial interactions, angiogenesis ^24^, apoptosis signalling ^25^, inflammation in obesity, metabolic syndrome ^26^, the study of hair cells of the inner ear ^27^ and even virulence in fungi ^28^ and parasites ^29^.

A stem cell study has shown the effect of certain growth factors on the differentiation and maturation of stem cells in organotypic cell lines. Growth factors secreted by mesenchymal stem cells control in situ endothelial cells, fibroblasts, and chondrocytes ^30^. Indirect co-cultivation of bone marrow-derived mesenchymal stromal cells and human umbilical vein endothelial cells in the “transwell system” leads to the formation of an optimally organized vascular structure ^31^. A system for co-culturing C2C12 and 3T3-L1 cells was created to understand the formation of muscle and fat cells in animals, which can be used to model muscle degeneration, apoptosis, and muscle regeneration ^32^. A triple co-culture system has been successfully created, including pig brain endothelial cells (PBEC), astrocytes and pericytes to create the blood-brain barrier model. It has been shown that primary pig astrocytes and pericytes are used for triple cell culture with PBEC ^33,34^.

Animal models are traditionally used to model cell transformation, EMT, and metastasis processes but have limitations due to the inability to take into account and control all experimental variables, as well as the insufficient resolution of the used analytical methods ^35^. Despite the fact that in vitro models provide a simplified view of the processes occurring during oncogenesis, they present a powerful tool that complements within in vivo research. This approach allows thorough analysis of molecular mechanisms under strictly controlled conditions by usage of human cells and ability to study single cells ^36^. Historically, the first in vitro cancer models were represented by two-dimensional cultures of immortalized cancer cell lines ^37^, but the role of the stromal cells in the development of cancer was not considered. However, researchers are increasingly attracted by the interaction between cancer cells and microenvironment, which regulate the behavior of the former ^38^. Thus, co-cultivation systems have been proposed for modeling interactions between cancer and microenvironment, ranging from two-dimensional, indirect co-cultures ^39^ to more modern systems based on a complex three-dimensional environment in which multiple cell types are placed ^40^.

Cell co-culture systems are widely used and capable of reproducing the conditions of cell-cell interactions. At the same time, the main disadvantage of this system is the high price and the inability to reproduce them independently in the laboratory. In addition, such liners are created from materials (polycarbonates) that cannot fully replicate the native adhesive surface for cells. The aim of this study is to obtain a system of co-cultivation of cells from available reagents directly in the laboratory.

## 2 RESULTS AND DISCUSSION

### 2.1 Membrane characterisation

Protein membranes with magnetic nanoparticles, were made from bovine serum albumin (BSA) (Sigma). For the formation of membranes, chemical crosslinking of proteins was performed by using 1-(3-Dimethylaminopropyl)-3-ethylcarbodiimide hydrochloride (EDC). The choice of a crosslinking agents is due to their low toxicity, while glutaraldehyde was not used to crosslink proteins due to excessive toxicity.

The thickness of the membranes were established by embedding the samples in paraffin and observing them by a light microscope, and by using a scanning electron microscope, after critical point drying (CPD) of the membranes. The thickness of protein membranes were in the range from 100 to 300 micrometers, depending on the volume of filling the wells in silicone form. The filling volume of 100 microliters corresponds to a thickness of 150 micrometers and a membrane area of 450 mm^2^. Membranes with a thickness of less than 100 microns are breakable and during polymerization in silicone form does not always completely cover the surface of the pit, and therefore may contain holes. On the other hand, It makes no sense to produce membranes with a thickness of 300 or more micrometers as their thickening does not give them any new properties for co-cultivation.

The membrane surface is determined by the relief of the embedding form, therefore, the modification of the latter leads to the change of the protein membrane microrelief. Membranes obtained by FDM and MSLA printers differ significantly in terms of their topography. On membranes obtained using an FDM printer, there is a pronounced relief of strips 0.4 mm wide corresponding to the diameter of the extruder used for printing (1 A). In this case, the height difference between the upper part of the relief and the lower is about 0.1 mm. In addition, due to imperfections in the printer’s feed system, the surface is rather heterogeneous. While the membranes obtained by using an MSLA printer appear to be much smoother. Withal, they also have a relief consisting of uniform 20 by 20 micrometer squares originating from the pixels of the LCD mask of the MSLA printer (1 B).

However, the membranes obtained by either method did not display any significant difference in terms of cell cultivation. This likely happened, due to the fact that used cell cultures were nonsensitive to the microrelief of the substrate. Although, the ability to create a relief and control it opens up opportunities for experiments with such cells (for example, cells of neurons, muscles).

### 2.2 Cells on membrane

Cells were grown on the protein membranes with and without nanoparticles, but no dependence of cell growth on the presence or absence of iron-carbon nanoparticles in the composition of the membranes was found. The abundance of nanoparticles required for levitation of a protein membrane in a culture medium under a constant magnetic field was 10 *µ*g per membrane. The observed difference between cell growth on membranes created using MSLA and FDM technologies was not found (2 A and B). However, it is worth noting that the height difference on the membranes made using FDM is quite large, which greatly complicates the work with cells on them using optical microscopy. As a result, all further experiments were carried out on membranes made using MSLA technology.

It was found that cells are able to form a monolayer on the membrane within the three days. Treating the surface of the membrane with collagen (500 *µ*g / ml), fibronectin (20 *µ*g/ml) or polylysine (100 *µ*g / ml) significantly improves the attachment of cells to the membrane and increase their growth rate. During growth, cells are able to form several layers on the membrane. In SEM, the formation of numerous cell processes (lamellipodia and filopodia) is visible, which indicates migration ability on the surface. However, after the monolayer is formed, cells stop their migration across the membrane. This was shown by culturing cells on a membrane with magnetic nanoparticles in the presence of a magnetic field in the volume of the culture medium with a clean bottom of the culture vessel. Within three days, no migration of cells from the membrane to the bottom of the culture vessel was detected (Figure 4).

**FIGURE 1.**
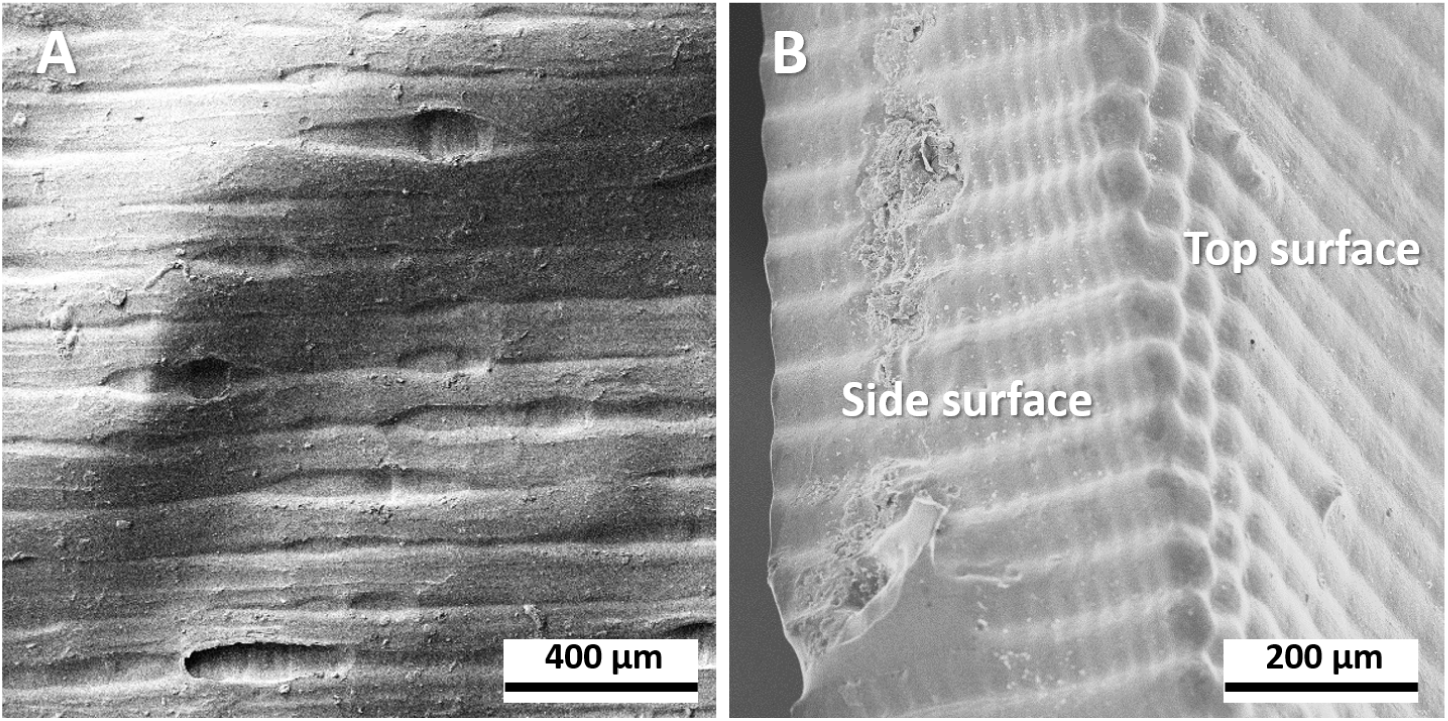
SEM microscopy images of FDM based membrane (A) and MSLA based membrane (B)

**FIGURE 2.**
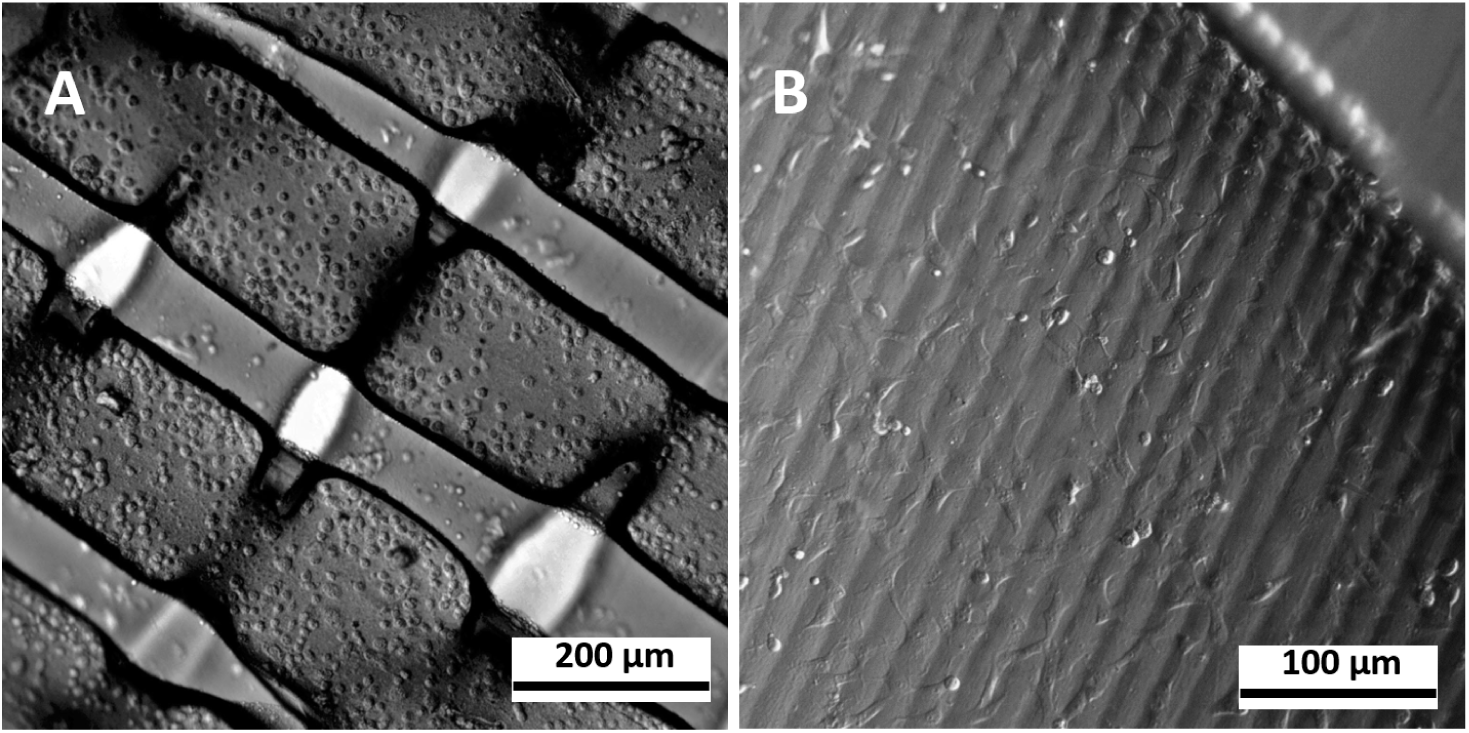
Optical microscopy of cells growing onto a surface of membranes, which was prepared in FDM (A) and MSLA (B) molds

**FIGURE 3.**
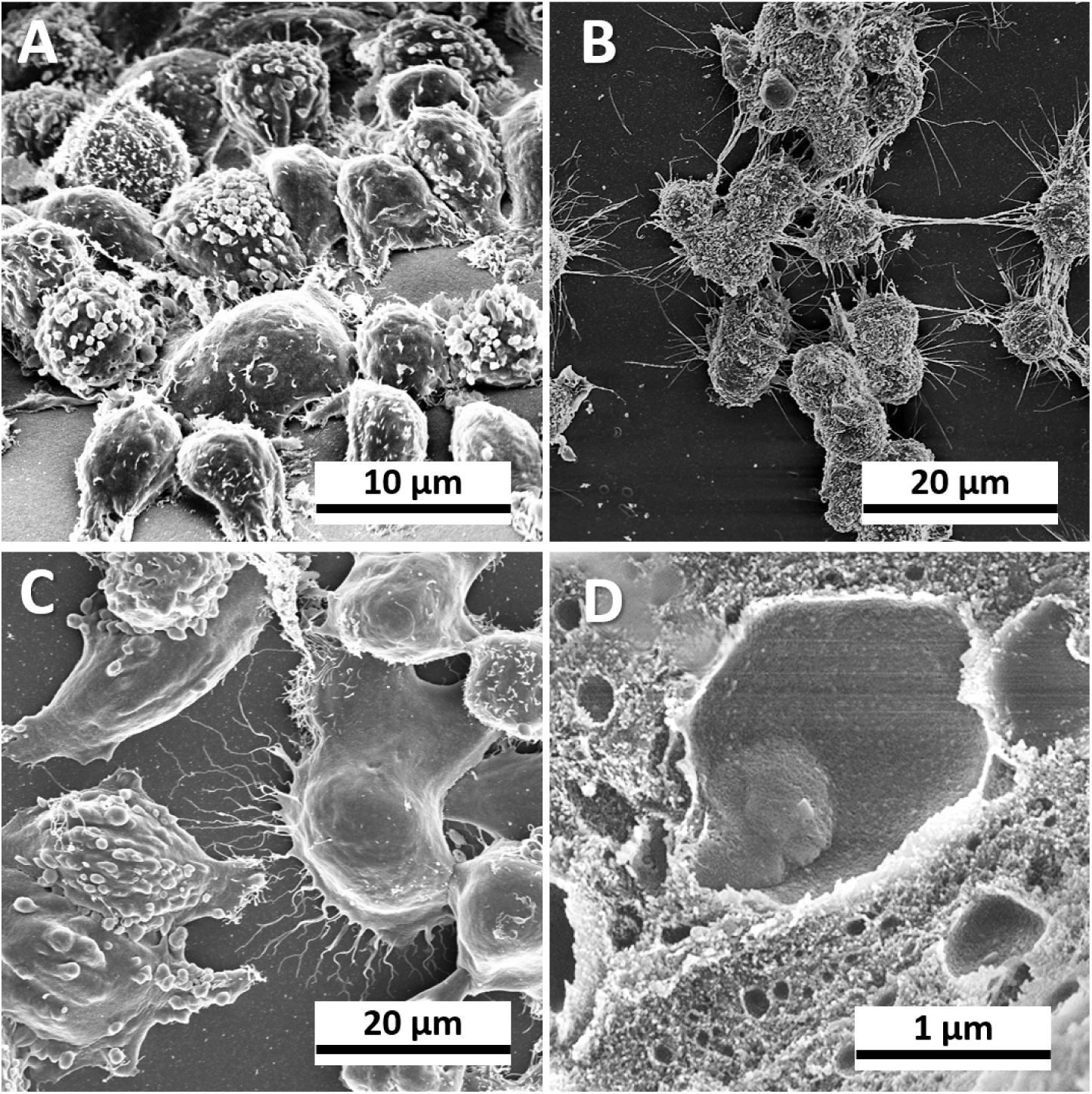
A, B MiaPaca cells on protein membrane. SEM images. C. HeLa cells on the protein membrane. D. Pores in the inside of the protein membrane. SEM images

**FIGURE 4.**
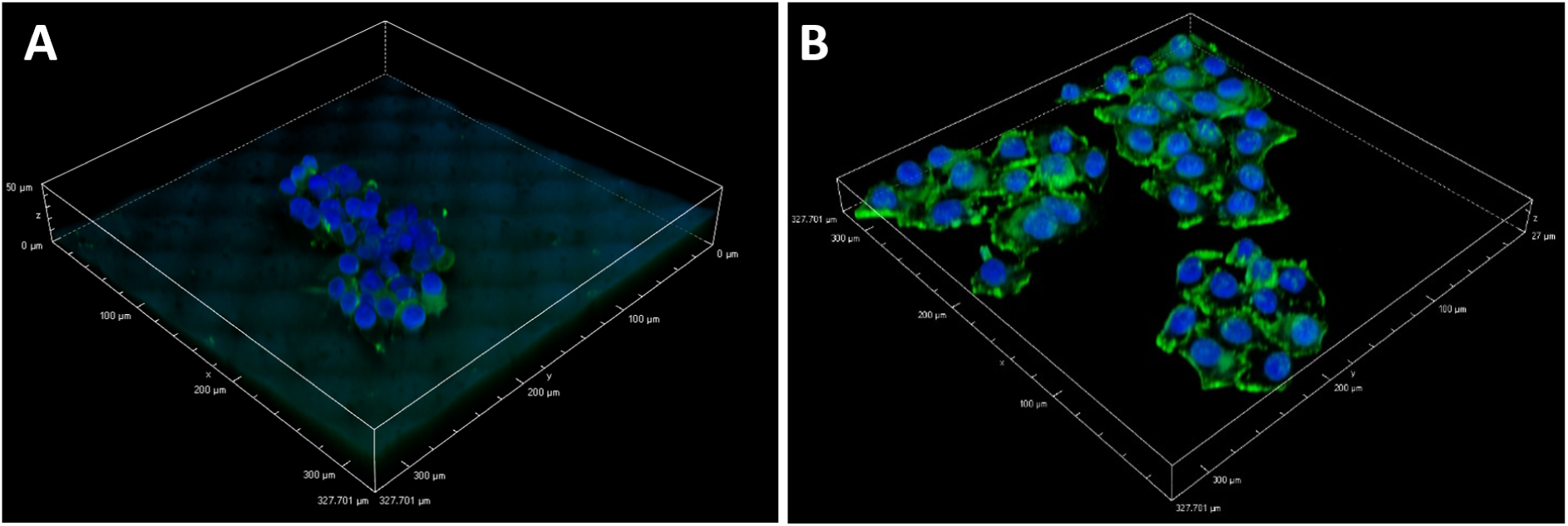
3d reconstructed image obtained with confocal microscopy of samples of cells on a membrane (A) and in dish(B).

The created membranes are permeable to low molecular weight substances and are biocompatible. By using the CPD, the structure and volume of protein membranes and pores inside it are preserved. Implication of SEM allowed observation of numerous variously sized pores on the cleaved membrane (Fig. 3). When membranes are placed in the culture medium, the pores inside them are filled with culture medium and factors secreted by the cells, the membrane swells slightly. Growth factors can infiltrate into such pores, followed by their time-dependent release. It was found that the membrane surface is inert to the cells and enzymes, and trypsin treatment to remove cells from the surface does not destroy protein membranes. These membranes are thin, permeable to metabolic products and growth factors, biocompatible.

Protein membranes were embedded in paraffin and resin (for electron microscopy), sectioned and stained (Fig. 5). Single cells and multilayer cell formations are visible on membrane samples. Intercellular contacts are intact, contact with the protein membrane is not destroyed. We have shown the ability to study cells on sections of the protein membrane, and analyze their structure without removing from the surface. The commercial analogs of cell co-cultivation systems do not provide for the possibility of obtaining quality membrane sections with cells. It was proposed analysis of cellular barrier models by Transmission Electron Microscopy (TEM), with transwell assays and demonstrate the ability of the method for assessment of nanoparticle transport across the Transwell membrane ^41^. At the same time, the obtained sections and images are of poor quality due to the membrane material. The protein membrane is much simpler and better for TEM studies. Making it possible to investigate the paracrine effect of cells on the formation of intercellular contacts and intracellular structures in situ without destruction of the integrity of intercellular contacts. In addition, the ability to detach cells from the membrane surface by using trypsin as in a method similar to detaching cells from plastic has been shown.

**FIGURE 5.**
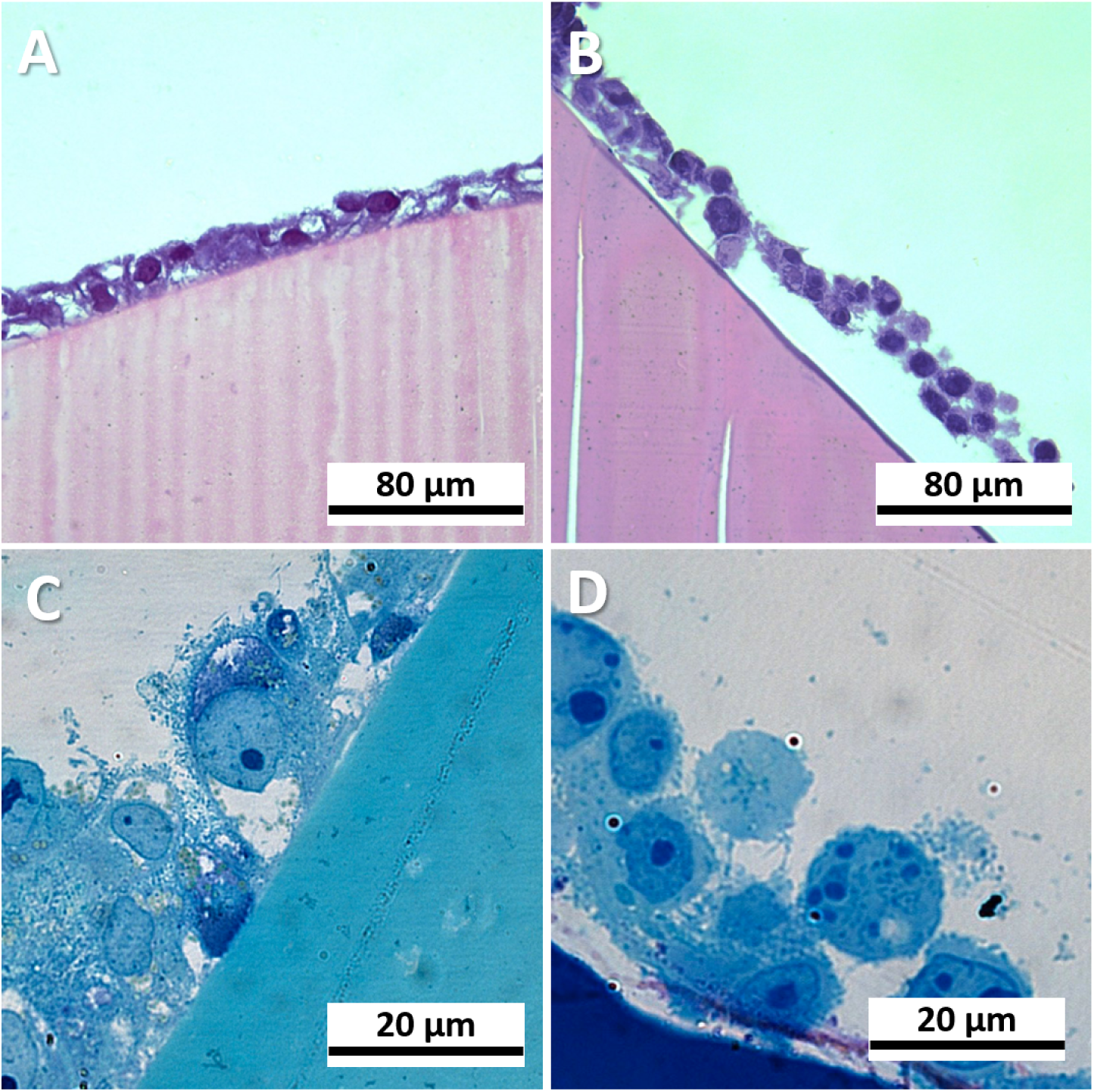
Light microscopy of cell cultures on proteins membranes. light microscopy of cell population on protein membrane (A). Paraffin section with HeLa cells (B) and MiaPaca2 cells (C). Semithin sections with HeLa cells (D) and MiaPaca2 cells (E).

To describe the changes in phenotypic cell markers during growth on a protein membrane, we studied expression of the cell markers characteristic for growth on a plastic surface (such as CK5.6, E-cadherin, vimentin, chromogranin A and synaptophysin but not CD56^42^. For studying the functional state we analyzed proliferative (Cyclin E2) and proapoptotic activity (p53, Bcl-2, annexin). It was found that with cell growth on the protein membrane, the expression profile of specific markers does not change.

## 3 CONCLUSION

The proposed cell culturing system could be manufactured within the laboratory conditions and present a cost-efficient alternative to the commercially available membrane inserts. Applying adhesive coatings (fibrillar proteins such as collagen and fibronectin) onto the protein membrane allows the cells to adhere to the bottom and maintain their viability.

We showed the possibility to study cells on sections of the protein membrane, to analyze their structure without removal from the surface. While, the commercial analogs of cell co-cultivation systems do not provide for the possibility of obtaining membrane sections. Thus, it becomes possible to investigate the paracrine effect of cells on the formation of intercellular contacts and intracellular structures.

## 4 MATERIALS AND METHODS

### 4.1 Synthesis and modification of nanoparticles

Metal-carbon NPs Fe@C were synthesized using the gas-condensation technique as described ^43^, in the applied magnetism laboratory of the Institute of Metal Physics, Ural Division of the Russian Academy of Sciences. A high-frequency alternating magnetic field overheated the iron drop to a temperature of about 2100°C in the flow of inert gas containing a hydrocarbon precursor. This leads to gradual evaporation of the molten iron drop. Hydrocarbon was catalytically decomposed on flying MNPs, forming a carbon shell covering the iron core [19]. Further, the NPs were cooled and collected by a textile filter. A chemical procedure using aryl-diazonium salts was applied to modify the surface of metal-carbon NPs Fe@C ^44,45^. To synthesize diazonium salts 0.1M sodium nitrite solution was added dropwise at 0-2°C in an ice bath not allowing overheating of the solution to 2 ml of 0.05M solution of 4-aminobenzylamine in 0.1 M hydrochloric acid until the positive reaction on starch-iodide paper occurred. The obtained solution of diazonium salts was added to the aqueous suspension of metal-carbon particles (100-500 mg) with ultrasonic treatment (22 kHz). After addition, the MNPs suspension was further sonicated for 15 minutes. The obtained MNPs were precipitated on a permanent magnet and washed three times with water and ethyl alcohol using magnetic decantation to eliminate reagents, then redispersed in deionized water.

Nanoparticles have a high magnetic moment of about 100 emu/g, which is more than that of iron oxide nanoparticles (60-70 emu/g) - this allows a more effective concentration of the membrane with a magnetic field. These nanoparticles, due to the carbon shell, are highly stable and do not degrade over time, in addition, they are biologically inert.

### 4.2 Magnetic system

A magnetic system for magnetically controlled cell co-cultivation was constructed based on commercially available permanent NiFeB magnets. The use of permanent magnets allows creation of a non-volatile system, so they were given preference to a system based on electromagnets.

The system was constructed of a cylindrical permanent magnet (d = 50 mm, h = 20 mm), a cylinder of the same size made of soft magnetic metal, which allows improving the uniformity of the magnetic field to increase the field above the magnet and another cylindrical permanent magnet (d = 10 mm, h = 3 mm), which allows to obtain the optimal configuration of the magnetic field, with the highest magnetic field strength in the projection of the center of the system. An additional central magnet not only increases the field and the gradient in Z (the field is 1.5 times, and the gradient is 3 times), but it also creates a much larger gradient in the radial component. All this allows to create the optimal configuration of the magnetic field for holding a magnetically controlled membrane at the phase boundary and, at the same time, its exact positioning in the center of the Petri dish. The magnetic system is placed inside the case printed on a 3D printer made of PLA plastic, equipped with interchangeable supports of different heights, allowing to adjust the distance to the Petri dish.

### 4.3 3d printing of primary mold

To create the primary molds, 3D printers based on different principles were used to determine which one is the most promising for creating objects with controlled shape and microrelief. We used FDM (fused deposition modeling) 3d printer CR-10s (Creality 3d, China) and MSLA (masked stereolithography) 3d printers Photon (Anycubic, China) and Mars (Elegoo, China). The FDM 3d printer was additionally modified using BL-touch, MSLA printers were used without any additional modifications. The printer software has been updated to the latest stable version. When preparing 3d models for printing on an FDM printer, the latest stable version of Prusa Slicer was used; for MSLA, the latest stable version of Chitubox was used. Printing profiles and 3d models in STL format are attached to supporting materials. Primary molds were printed on an FDM 3d printer out of PLA plastic (Bestfilament, Russia), on MSLA 3d printers out of SLA resin (Anycubic, China).

For the manufacture of secondary molds, two-component Alcorsil 315 silicone (Alcorsil, China) was added to the primary ones. A minimally viscous silicone was used, so there was no need for evacuation to remove bubbles from the silicone mass. After hardening, the secondary molds were washed in 99% isopropyl alcohol to remove any impurities.

### 4.4 Manufacturing of protein membranes

Protein, aminated nanoparticles, and crosslinkers were placed in silicone forms and chemical crosslinking of magnetic nanoparticles with proteins occurred. As a result a membrane with a thickness of 100-150 *µ*m was obtained.

The choice of bovine serum albumin as a main component for the membrane is due to its properties such as biodegradability, accessibility for any laboratory, and good intermolecular cross-linking ability. BSA 20% solutions in deionized water are crosslinked to the functional groups of aminate nanoparticles with EDC crosslinker (1-(3-Dimethylaminopropyl)-3-ethylcarbodiimide hydrochloride). 25 *µ*l of iron-carbon nanoparticles, grafted with aminogroups (4 mg/ml) is added to 1 ml of the solution BSA and sterilized through a 0.22 *µ*m filter. EDC-BSA crosslinking was carried out in a 0.05 M EDC in a ratio of 1:1 for 20 minutes at room temperature.

Protein membranes were rewashed by sterile deionized water, glutamine (for three times), and then incubated overnight in a full culture medium. Before seeding cells on the membrane, it was washed with culture medium for several times. Components of the extracellular matrix (collagen, fibronectin) were applied to the surface of the membrane to create adhesive properties of the membrane. The obtained membranes are durable, do not stick together and could be straightened by constant magnetic field.

### 4.5 The cell cultures

MiaCaPa-2 tumor cell lines, HEK, HeLa and MRC-5 fetal lung fibroblasts were obtained from the Russian cell cultures collection of the Institute of Cytology RAS (Russia, St. Petersburg). The cell culture was maintained in plastic vials containing culture medium (DMEM) with 10% FBS and 50 *µ*g / ml gentamicin, in an incubator at + 37 °C, 5% CO2. A magnetic holder is placed on top of a Petri dish with cell cultures.

### 4.6 Cell viability assessment

MTT assay was carried out for assessing cell metabolic activity. 50 *µ*L of serum-free media and 50 *µ*L of MTT solution at 4 mg/ml in PBS to 100*µ*l of media was added into each well (37°C for 3 hours). After incubation, added 150 *µ*L of MTT solvent (4 mM HCl, 0.1% NP40 in isopropanol) into each well (incubating for 15 minutes). Plate was read within 1 hour at OD=590 nm ^46^ using a plate reader.

### 4.7 Histology, immunocytochemistry and electron microscopy

For cell sections the protein membranes were fixed in 4% paraformaldehyde, dehydrated in alcohol and xylene, and embedded in paraffin using the same method as tissue sample processing. The resulting cell blocks were trimmed and sectioned at 1 *µ*m with a rotary microtome. The sections were stained with Hematoxylin-Eosin for general morphology. Immunocytochemical investigation of the cell culture’s functional state was carried out by using a Nikon confocal microscope with a motorized table and specific antibodies.

Cells with membranes for SEM were fixed in 2% paraformaldehyde and 2.5% glutaraldehyde in cacodylate buffer ^47^ with 5% sucrose, then fixed in 2% osmium tetroxide. The samples were dehydrated in alcohol and acetone, then dried to a critical point in a K850 Critical Point Dryer (Quorum Technologies). A 15 nm gold/palladium layer was deposited on the surface of the samples in a Q150T Plus metal sputtering device (Quorum Technologies). Furtherly, cells and protein membranes were investigated in an AURIGA FIB-SEM workstation scanning electron microscope («Carl Zeiss; MT», Oberkochen, Germany) in SE detector in the range of magnifications 50-50000. To visualize cell contact with a protein membrane in the cross section, protein membranes were fixed in 2% paraformaldehyde and 2.5% glutaraldehyde on cacodylate buffer pH 7.2 (Karnovsky, 1965) with 5% sucrose, then post-fixed in 2% osmium tetroxide, contrasted with uranyl acetate en bloc. The material was dehydrated in alcohol and acetone, then embedded in epoxy resin (Spurr). Semithin (900 nm thick) sections of epoxy blocks were stained with toluidine blue with the addition of 1% borax and examined under an optical microscope ^48^.

### 4.8 Confocal microscopy examination

Permeabilization of the samples was carried out with a 0.5% Triton X-100 solution for 15 minutes, followed by three times washing with PBS for 5 minutes each.

Samples were stained with ReadyProbes ActinGreen 488 and ReadyFlow Hoechst 33342 (Thermo Fisher Scientific, USA) according to the manufacturer’s recommendations.

The A1R confocal laser scanning microscope based on the ECLIPSE Ti-E inverted microscope equipped with the CFI Plan Fluor 20XC MI multi-immersion lens (Nikon, Japan) was used to obtain images. The photography was carried out in water immersion. The excitation / emission wavelengths for Hoechst 33342 were: ex = 405 nm / em = 450 (50) nm, for AlexaFluor 488: ex = 488 nm / em = 525 (50) nm. The collection and processing of images was carried out in the program NIS Elements (ver. 4.50) (Nikon, Japan)

## 4.9 Acknowledgements

The reported study was funded by the Russian Science Foundation Grant #19-74-00081

## References

1. Renaud J, Martinoli MG. Development of an insert co-culture system of two cellular types in the absence of cell-cell contact. JoVE (Journal of Visualized Experiments) 2016(113): e54356.

2. Hatherell K, Couraud PO, Romero IA, Weksler B, Pilkington GJ. Development of a three-dimensional, all-human in vitro model of the blood–brain barrier using mono-, co-, and tri-cultivation Transwell models. Journal of neuroscience methods 2011; 199(2): 223–229.

3. Grobstein C. Morphogenetic interaction between embryonic mouse tissues separated by a membrane filter. Nature 1953; 172(4384): 869–871.

4. Lander AD. How cells know where they are. Science 2013; 339(6122): 923–927.

5. Kelso A. Cytokines: principles and prospects. Immunology and cell biology 1998; 76(4): 300–317.

6. Nitta CF, Orlando RA. Crosstalk between immune cells and adipocytes requires both paracrine factors and cell contact to modify cytokine secretion. PLoS One 2013; 8(10).

7. Talayev VY, Matveichev A, Lomunova M, et al. The effect of human placenta cytotrophoblast cells on the maturation and T cell stimulating ability of dendritic cells in vitro. Clinical & Experimental Immunology 2010; 162(1): 91–99.

8. Elishmereni M, Alenius H, Bradding P, et al. Physical interactions between mast cells and eosinophils: a novel mechanism enhancing eosinophil survival in vitro. Allergy 2011; 66(3): 376–385.

9. Dono M, Zupo S, Massara R, et al. In vitro stimulation of human tonsillar subepithelial B cells: requirement for interaction with activated T cells. European journal of immunology 2001; 31(3): 752–756.

10. Bournival J, Plouffe M, Renaud J, Provencher C, Martinoli MG. Quercetin and sesamin protect dopaminergic cells from MPP. Oxidative medicine and cellular longevity 2012; 2012.

11. Zhu L, Bi W, Lu D, Zhang C, Shu X, Lu D. Luteolin inhibits SH-SY5Y cell apoptosis through suppression of the nuclear transcription factor-,*c*B, mitogen-activated protein kinase and protein kinase B pathways in lipopolysaccharide-stimulated cocultured BV2 cells. Experimental and therapeutic medicine 2014; 7(5): 1065–1070.

12. Kim JY, Kim DH, Kim JH, et al. Umbilical cord blood mesenchymal stem cells protect amyloid-*/]*42 neurotoxicity via paracrine. World journal of stem cells 2012; 4(11): 110.

13. Mauri M, Lentini D, Gravati M, et al. Mesenchymal stem cells enhance GABAergic transmission in co-cultured hippocampal neurons. Molecular and Cellular Neuroscience 2012; 49(4): 395–405.

14. Iwashita M, Sakoda H, Kushiyama A, et al. Valsartan, independently of AT1 receptor or PPAR*y*, suppresses LPS-induced macrophage activation and improves insulin resistance in cocultured adipocytes. American Journal of Physiology-Endocrinology and Metabolism 2012; 302(3): E286–E296.

15. De Boer AA, Monk JM, Robinson LE. Docosahexaenoic acid decreases pro-inflammatory mediators in an in vitro murine adipocyte macrophage co-culture model. PloS one 2014; 9(1).

16. Lawrenson K, Grun B, Benjamin E, Jacobs IJ, Dafou D, Gayther SA. Senescent fibroblasts promote neoplastic transformation of partially transformed ovarian epithelial cells in a three-dimensional model of early stage ovarian cancer. Neoplasia (New York, NY) 2010; 12(4): 317.

17. Liu J, Joha S, Idziorek T, et al. BCR-ABL mutants spread resistance to non-mutated cells through a paracrine mechanism. Leukemia 2008; 22(4): 791–799.

18. Gupta D, Treon S, Shima Y, et al. Adherence of multiple myeloma cells to bone marrow stromal cells upregulates vascular endothelial growth factor secretion: therapeutic applications. Leukemia 2001; 15(12): 1950–1961.

19. Karadag A, Oyajobi BO, Apperley JF, Russell R, Croucher PI. Human myeloma cells promote the production of interleukin 6 by primary human osteoblasts.. British journal of haematology 2000; 108(2): 383–390.

20. Moore MB, Kurago ZB, Fullenkamp CA, Lutz CT. Squamous cell carcinoma cells differentially stimulate NK cell effector functions: the role of IL-18. Cancer Immunology, Immunotherapy 2003; 52(2): 107–115.

21. Zhang M, Xu Mx, Zhou Z, et al. The differentiation of human adipose-derived stem cells towards a urothelium-like phenotype in vitro and the dynamic temporal changes of related cytokines by both paracrine and autocrine signal regulation. PloS one 2014; 9(4).

22. Qu C, Puttonen KA, Lindeberg H, et al. Chondrogenic differentiation of human pluripotent stem cells in chondrocyte co-culture. The international journal of biochemistry & cell biology 2013; 45(8): 1802–1812.

23. Ichikawa J, Okada A, Taguchi K, et al. Increased crystal–cell interaction in vitro under co-culture of renal tubular cells and adipocytes by in vitro co-culture paracrine systems simulating metabolic syndrome. Urolithiasis 2014; 42(1): 17–28.

24. Beckner ME, Jagannathan S, Peterson VA. Extracellular angio-associated migratory cell protein plays a positive role in angiogenesis and is regulated by astrocytes in coculture. Microvascular research 2002; 63(3): 259–269.

25. Vjetrovic J, Shankaranarayanan P, Mendoza-Parra MA, Gronemeyer H. Senescence-secreted factors activate M yc and sensitize pretransformed cells to TRAIL-induced apoptosis. Aging cell 2014; 13(3): 487–496.

26. Suganami T, Nishida J, Ogawa Y. A paracrine loop between adipocytes and macrophages aggravates inflammatory changes: role of free fatty acids and tumor necrosis factor *α*. Arteriosclerosis, thrombosis, and vascular biology 2005; 25(10): 2062–2068.

27. May LA, Kramarenko II, Brandon CS, et al. Inner ear supporting cells protect hair cells by secreting HSP70. The Journal of clinical investigation 2013; 123(8): 3577–3587.

28. Dagenais TR, Giles SS, Aimanianda V, Latgé JP, Hull CM, Keller NP. Aspergillus fumigatus LaeA-mediated phagocytosis is associated with a decreased hydrophobin layer. Infection and immunity 2010; 78(2): 823–829.

29. Spiliotis M, Lechner S, Tappe D, Scheller C, Krohne G, Brehm K. Transient transfection of Echinococcus multilocularis primary cells and complete in vitro regeneration of metacestode vesicles. International journal for parasitology 2008; 38(8-9): 1025–1039.

30. Wang Y, Chen X, Cao W, Shi Y. Plasticity of mesenchymal stem cells in immunomodulation: pathological and therapeutic implications. Nature immunology 2014; 15(11): 1009.

31. Böhrnsen F, Schliephake H. Supportive angiogenic and osteogenic differentiation of mesenchymal stromal cells and endothelial cells in monolayer and co-cultures. International journal of oral science 2016; 8(4): 223–230.

32. Pandurangan M, Hwang I. Application of cell co-culture system to study fat and muscle cells. Applied microbiology and biotechnology 2014; 98(17): 7359–7364.

33. Thomsen LB, Burkhart A, Moos T. A triple culture model of the blood-brain barrier using porcine brain endothelial cells, astrocytes and pericytes. PloS one 2015; 10(8).

34. Stone NL, England TJ, O’Sullivan SE. A Novel Transwell Blood Brain Barrier Model Using Primary Human Cells. Frontiers in Cellular Neuroscience 2019; 13: 230. doi: 10.3389/fncel.2019.00230

35. Evans JP, Sutton PA, Winiarski BK, et al. From mice to men: Murine models of colorectal cancer for use in translational research. Critical reviews in oncology/hematology 2016; 98: 94–105.

36. Shologu N, Szegezdi E, Lowery A, Kerin M, Pandit A, Zeugolis DI. Recreating complex pathophysiologies in vitro with extracellular matrix surrogates for anticancer therapeutics screening. Drug discovery today 2016; 21(9): 1521–1531.

37. Sanford KK, Barker BE, Woods MW, Parshad R, Law LW. Search for “indicators” of neoplastic conversion in vitro. Journal of the National Cancer Institute 1967; 39(4): 705–733.

38. Junttila MR, Sauvage dFJ. Influence of tumour micro-environment heterogeneity on therapeutic response. Nature 2013; 501(7467): 346–354.

39. Katt ME, Placone AL, Wong AD, Xu ZS, Searson PC. In vitro tumor models: advantages, disadvantages, variables, and selecting the right platform. Frontiers in bioengineering and biotechnology 2016; 4: 12.

40. Bersini S, Jeon JS, Moretti M, Kamm R. In vitro models of the metastatic cascade: from local invasion to extravasation. Drug discovery today 2014; 19(6): 735–742.

41. Ye D, Dawson KA, Lynch I. A TEM protocol for quality assurance of in vitro cellular barrier models and its application to the assessment of nanoparticle transport mechanisms across barriers. Analyst 2015; 140: 83–97. doi: 10.1039/C4AN01276C

42. Gradiz R, Silva HC, Carvalho L, Botelho MF, Mota-Pinto A. MIA PaCa-2 and PANC-1–pancreas ductal adenocarcinoma cell lines with neuroendocrine differentiation and somatostatin receptors. Scientific reports 2016; 6: 21648.

43. Yermakov A, Uimin M, Byzov I, et al. Structure and magnetic properties of carbon encapsulated FeCo@C and FeNi@C nanoparticles. Materials Letters 2019; 254: 202–205. doi: 10.1016/j.matlet.2019.07.067

44. Postnikov PS, Trusova ME, Fedushchak Ta, Uimin Ma, Ermakov aE, Filimonov VD. Aryldiazonium tosylates as new efficient agents for covalent grafting of aromatic groups on carbon coatings of metal nanoparticles. Nanotechnologies in Russia 2010; 5(7-8): 446–449. doi: 10.1134/S1995078010070037

45. Minin AS, Uymin MA, Yermakov AY, et al. Application of NMR for quantification of magnetic nanoparticles and development of paper-based assay. Journal of Physics: Conference Series 2019; 1389(November): 012069. doi: 10.1088/1742-6596/1389/1/012069

46. Tang XJ, Huang KM, Gui H, et al. Pluronic-based micelle encapsulation potentiates myricetin-induced cytotoxicity in human glioblastoma cells. International Journal of Nanomedicine 2016; 11: 4991–5002. doi: 10.2147/IJN.S114302

47. Karunovsky M. A formaldehyde-glutaraldehyde fixative of high osmolality for use in electron microscopy. J. Cell Biol. 1965; 27: 137A.

48. Payne JW. Polymerization of proteins with glutaraldehyde. Soluble molecular weight markers. Biochemical Journal 1973; 135(4): 867–873. doi: 10.1042/bj1350867

